# Cell-accurate optical mapping across the entire developing heart

**DOI:** 10.1101/143057

**Authors:** Michael Weber, Nico Scherf, Peter Kohl, Jan Huisken

**Affiliations:** Max Planck Institute of Molecular Cell Biology and Genetics, 01307 Dresden, Germany; Harvard Medical School, Boston, MA 02115, USA; Max Planck Institute for Human Cognitive and Brain Sciences, 04103 Leipzig, Germany; Institute for Experimental Cardiovascular Medicine, University Heart Centre Freiburg Bad Krozingen, School of Medicine, Albert-Ludwigs University, 79110 Freiburg, Germany; Morgridge Institute for Research, Madison, WI 53715, USA

## Abstract

Organogenesis depends on orchestrated interactions between individual cells and morphogenically relevant cues at the tissue level. This is true for the heart, whose function critically relies on well-ordered communication between neighbouring cells, which is established and fine-tuned during development. For an integrated understanding of the development of structure and function, we need to move from isolated snap-shot observations of either microscopic or macroscopic parameters to simultaneous and, ideally continuous, cell-to-organ scale imaging. We introduce cell-accurate three-dimensional Ca^2+^-mapping of all cells in the entire heart during the looping stage in live embryonic zebrafish, using high-speed light sheet microscopy and tailored image processing and analysis. We show how myocardial region-specific heterogeneity in cell function emerges during early development and how structural patterning goes hand-in-hand with functional maturation of the entire heart. Our method opens the way to systematic, scale-bridging, *in vivo* studies of vertebrate organogenesis by cell-accurate structure-function mapping across entire organs.

## Introduction

Organogenesis builds on cell-cell interactions that shape tissue properties, and tissue-level cues that control maturation of cell structure and function. During cardiogenesis, region-specific heterogeneity in inter-cellular activity patterns evolves as the heart undergoes large-scale morphological changes: The spontaneously active heart tube develops into the mature heart, in which pacemaker cells near the inflow site initiate the rhythmic excitation that spreads with differential velocities through distinct regions of the myocardium. This controlled cardiac activation gives rise to an orderly sequence of atrial and ventricular calcium release and contraction. An integrated understanding of cardiogenesis at the systems level requires simultaneous cell and organ scale imaging, under physiological conditions *in vivo.* Here, we present a high-speed light sheet microscopy and data analysis pipeline to measure fluorescent reporters of cardiomyocyte location and activity across the entire heart in living zebrafish embryos during the crucial looping period from 36 to 52 hours post fertilization (hpf). By noninvasively reconstructing the maturation process of the myocardium in its entirety at cellular resolution, our approach offers an integrative perspective on tissue and cell level simultaneously, which has previously required separate experimental setups and specimens. Our method opens a new way towards systematic assessment of the mutual interrelations between cell- and tissue properties during organogenesis.

## Results

The zebrafish is an appealing vertebrate model system with a simple, yet functionally conserved heart, and light sheet microscopy has proven to be supremely suited for obtaining *in vivo* recordings of the intact embryonic zebrafish heart^1,2^. Entire cardiac cycles were reconstructed in 4D (3D + time) using post-acquisition synchronization of high-speed light sheet movies in a z-stack. The resulting effective temporal resolution of about 400 volumes per second^3^ is unmatched by other *in vivo* volumetric imaging techniques such as light sheet microscopy with electrically focus-tunable lenses or swept, confocally-aligned planar excitation^4,5,6,7^. We built a light sheet microscope tailored for dual-color high-speed recording of cardiac activation in the living zebrafish embryo (Methods).

We investigated whether post-acquisition synchronization could be extended to visualizing calcium transients in cardiac myocytes across the entire heart of living embryonic zebrafish expressing the fluorescent calcium reporter GCaMP5G under the myl7 promoter (Fig. 1a, Supplementary Fig. 1). The genetically expressed calcium reporter provided a specific, consistent and non-invasive readout of cardiomyocyte activity *in vivo* (Fig. 1b, Videos 1 and 2). In a side-by-side comparison, the calcium signal had good and stable fluorescent yield at low excitation power, superior to genetically expressed voltage reporters. Importantly, the calcium signal faithfully reported presence and timing of cell activation (Supplementary Fig. 2)^8^. To prevent interference from tissue movement and deformation with observed signals, we decoupled electrical excitation and mechanical contraction by inhibiting the formation of the calcium-sensitive regulatory complex within sarcomeres, using a morpholino against tnnt2a (Methods). By mounting zebrafish embryos in low concentration agarose inside polymer tubes, we could position the embryos for precise optical investigation without anesthesia (Supplementary Fig. 1a, b). To attribute calcium dynamics to individual cardiomyocytes, we also recorded a fluorescent nuclear marker (myl7:H2A-mCherry). The high temporal (400 Hz) and spatial sampling (0.5 μm pixel size) was adequate for computing normalized average calcium transients throughout the cardiac cycle for each cell across the entire heart (Supplementary Fig. 3, Video S3, Methods).

**Figure 1.**
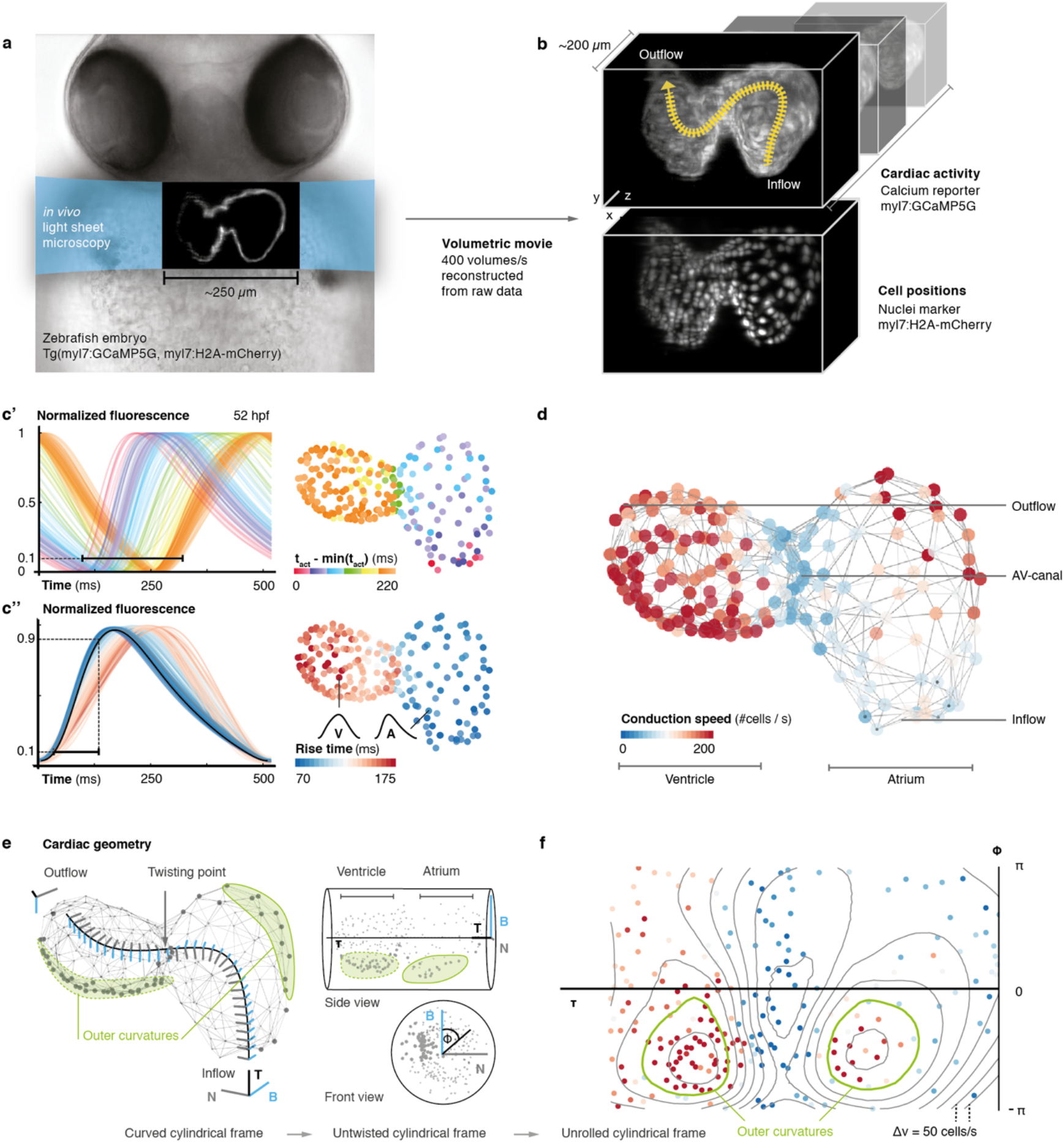
In vivo 3D optical mapping reveals cell-specific calcium transient patterns at 52 hours post fertilization (hpf). (**a**) Transmitted light microscopy image with ~250-μm-sized, two-chambered heart (shown as fluorescence image with light sheet illumination path). (**b**) Genetically encoded fluorescent markers expressed in myocardial cells report calcium transient activity and cell positions. Volumetric movies were reconstructed from multiple high-speed movies, each with a temporal resolution of 2.5 ms and a voxel size of 0.5 μm in *xy* and 1 μm in *z.* (**c**’) Normalized fluorescence plot of every cell’s calcium transient over one cardiac cycle. The network’s activation timing (t_act_) is visualized in 3D based on the time-point of 10% calcium transient amplitude in every individual cell (right, same color scale). (**c**”) Normalized fluorescence plot of all calcium transients, aligned in time based on the timing of deviation from minimal fluorescence intensity (3D network, same color scale). (**d**) Biological conduction speed, expressed as cells activated per unit of time, is visualized on the 3D network. (**e**) The basis vectors of the local coordinate system (tangent – black, normal – grey, and binormal – blue) are shown, moving along the centerline. The initial and final orientation of the moving reference frame is shown in a zoomed version at inflow and outflow sites. Outer curvature regions of atrium and ventricle are highlighted in green. (**f**) The unrolled cylinder results in a representation of 3D network function in a 2D map. Projections show conduction speed across the entire myocardium; iso-velocity lines (Δv = 50 cells/s) are shown in black. Outer curvature regions in atrium and ventricle are highlighted in green.

**Figure 2.**
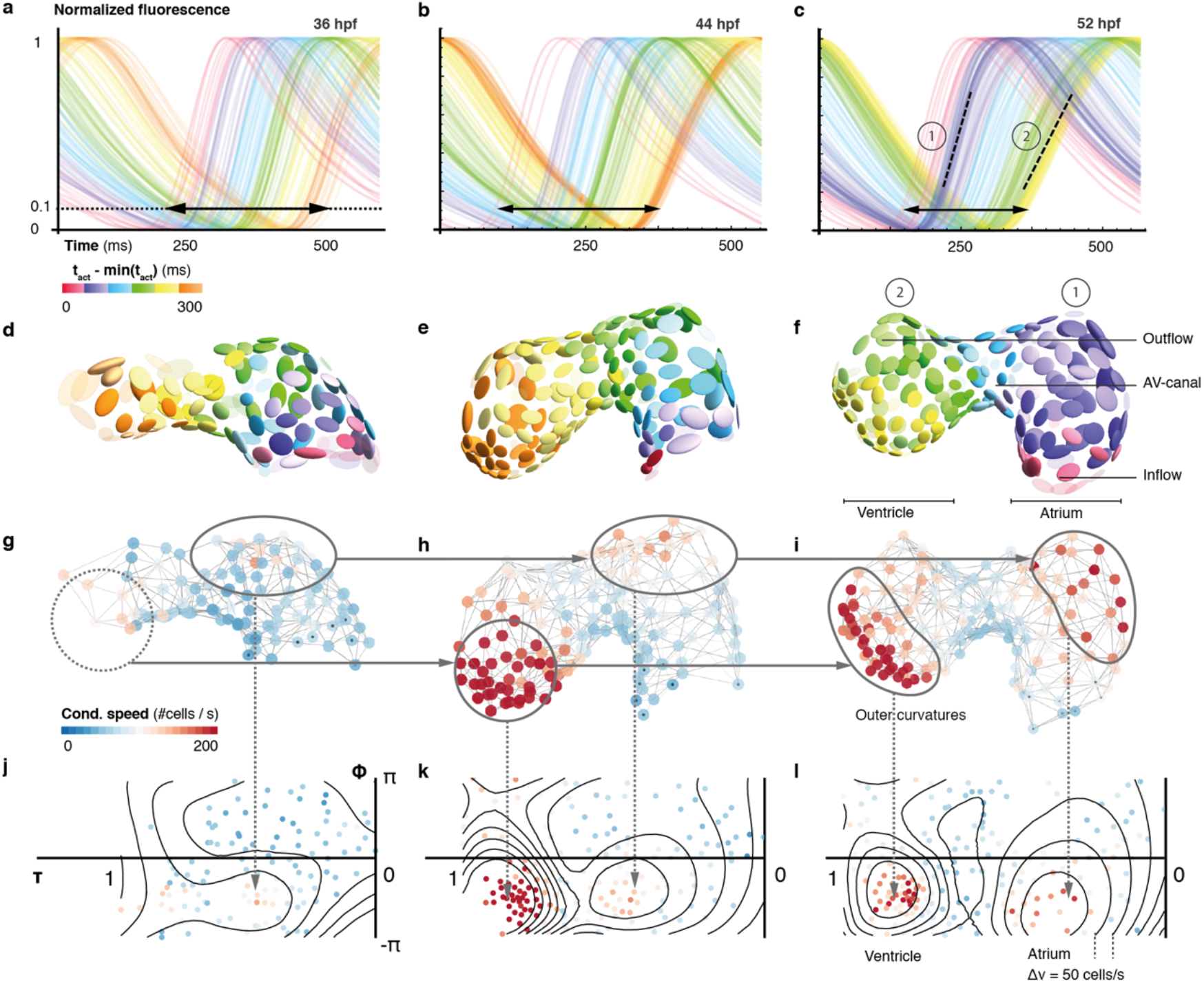
Organ maturation and functional cell remodelling from 36 to 52 hpf. (**a-c**) Normalized calcium transients of all cells are shown at three different time-points: (**a**) 36, (**b**) 44, and (**c**) 52 hpf. The shortening of black arrows indicates the decrease in time between earliest and latest activation of cells across the whole heart. The formation of atrial (1) and ventricular (2) populations with different calcium transients is highlighted by the dashed lines at 52 hpf. The color of each transient indicates its activation time (all three plots use the same color scale). (**d-f**) Changes in cardiac and cellular morphology at each developmental time point. Estimated cell shapes are visualized as scaled ellipsoids. The color of each ellipsoid indicates activation time; color code as in (a-c). Two main population of cells corresponding to highlighted clusters in (c) are indicated by numbers. (**g-i**) 3D pattern of biological conduction speeds across the heart. The formation of outer curvature clusters of cells over time is highlighted by arrows. (**j-l**) Development of conduction speeds across the myocardium, shown as 2D projections for the three time points. The color code indicates conduction speed. Isovelocity lines are shown in black (Δv = 50 cells/s). Corresponding outer curve cell populations are highlighted by arrows.

To map cellular activation timings onto a 3D structural representation of the heart, we identified every single cell’s activity during the cardiac cycle (Fig 1c’) and quantified dynamic characteristics of all cells across space and time. The distribution of calcium transient rise times (from 10% to 90% of calcium transient peak amplitude) revealed the emergence of distinct upstroke characteristics in different locations within the 3D network (by 52 hpf). Rise times were shortest for atrial muscle cells, intermediate in the atrioventricular canal (AVC), and longest for ventricular cardiomyoctes (Fig. 1c”), in keeping with higher vertebrates where atrial myocyte contraction is faster than that of ventricular cells^9^.

To get a more quantitative understanding of the 3D distribution of activity patterns, a canonical description of cell locations was needed. We traced and parameterized the myocardium’s centerline (Supplementary Fig. 4a, Methods), along which we assigned a unique position to each cell between inflow and outflow. We identified a positive correlation between rise time and position of cells along the midline, with a clear discontinuity at the AVC (Supplementary Fig. 4b), illustrating emergence of chamber-specific patterns of individual cell activation properties across the heart.

Next, we studied the spatial patterns of sequential cell activation, as a read-out for the speed of electrical conduction across the heart. While cardiac activation in the early linear heart tube is slow and uniform, the chambered heart shows areas of elevated conduction speed^1,10^. Common imaging-based methods for determining cell conduction speed are inherently biased: Either they are based on two-dimensional measurements and cannot faithfully quantify conduction speed due to loss of depth information, or they cannot resolve differences in local cell sizes and therefore only deliver the metric, or ‘biophysical speed’ of conduction (distance over time). To reflect biological progression of activation between cells (of potentially different size), we assessed local cell topology across the entire heart (Fig. 1d, Supplementary Fig. 5, Methods) to also calculate the ‘biological speed’ (number of cell over time) of conduction (Video S4). Our analysis revealed that biophysical and biological speed of conduction show differences between and within anatomical regions of the heart (Fig. 3). Either descriptor shows particularly slow conduction between the most proximal atrial cells and between the cells of the AVC, and faster conduction among working myocardial cells of the atrium and ventricle. In the atrium, there is a bias towards higher biophysical speeds, due to larger cell dimensions (Fig. 1d, Supplementary Fig. 4c).

**Figure 3.**
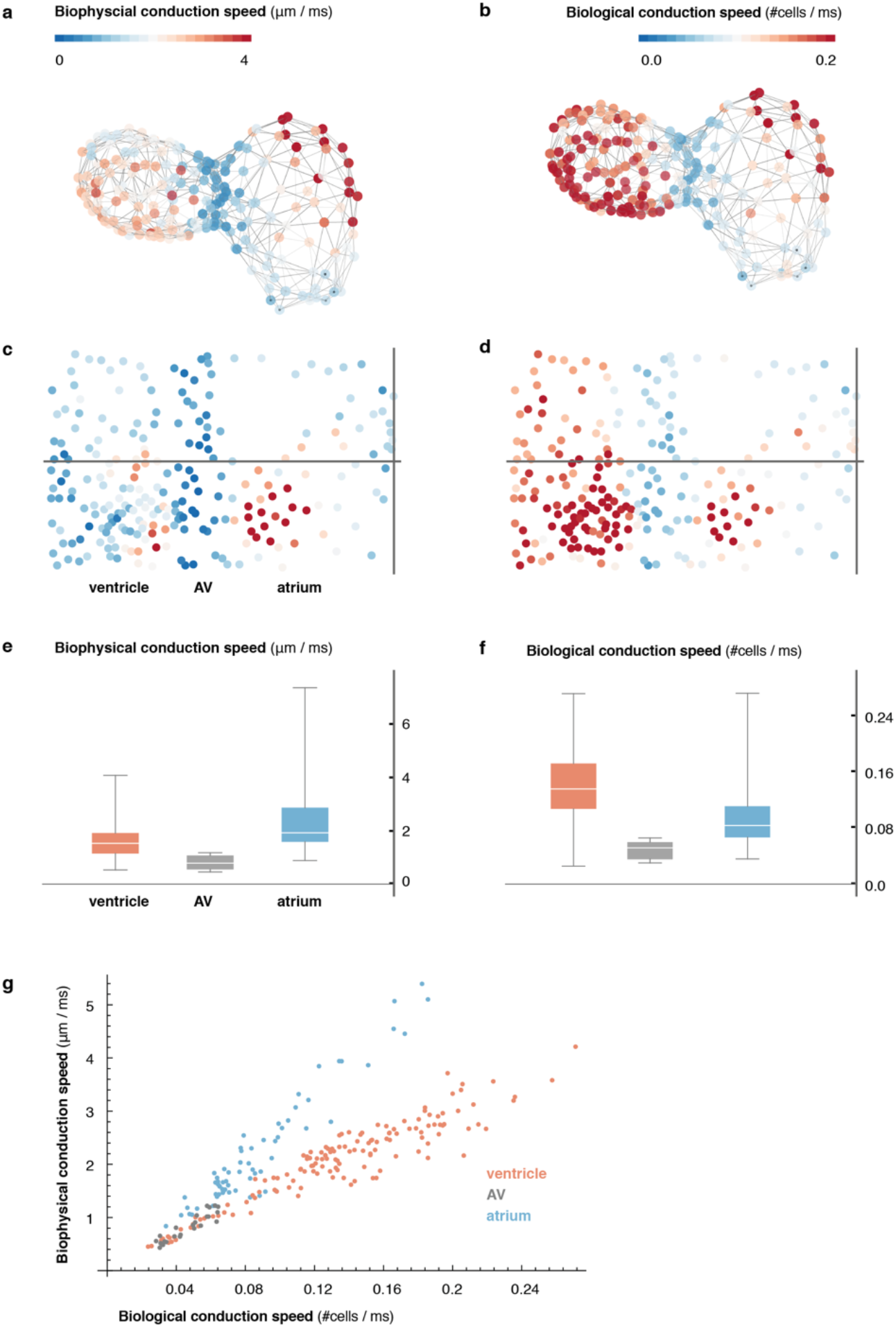
Comparison of metric and cell-to-cell speed measurements. (**a**) Biophysical conduction speed, expressed in μm / ms, is visualized on the 3D network. (**b**) Visualization of biological conduction speed, expressed as cells activated per unit of time on the same topology as (a). (**c, d**) Projection show biophysical (c) and biological (d) conduction speed across the entire myocardium. Same color code as in (a, b). (**e, f**) Descriptive statistics of conduction speeds in (manually defined) regions corresponding to atrial (blue), AVC (gray), and ventricular part (orange) of myocardium for biophysical (e) and biological (f) conduction speed. (**g**) Scatterplot showing chamber-specific differences in correlation of biological (x-axis) vs. biophysical (y-axis) conduction speed for atrial (blue), AVC (gray) and ventricular (orange) cells.

The cellular activation sequence in the pacemaker region is of particular importance to heart physiology and function, yet individual cells that give rise to earliest activation were difficult to identify with previous methods^11^. We found that, at 52 hpf, less than 10 cells per heart serve as activation origins. They are located in the sinus venosus at the heart’s inflow, which is a homologue of the cardiac pacemaker region in adult heart^12^. Our data further shows that the ring-like arrangement of pacemaker cells (Supplementary Fig. 6), together with a preferential orientation of myocardial cells in that region perpendicular to the inflow-outflow direction^13^, generates the initial planar activation front that propagates into the atrium.

To visualize the 3D conduction pattern, the heart can be represented as a curved cylinder with varying diameter and s-shaped deformations in the coronal and in the sagittal plane. We used the position along the midline (τ) and the local Frenet-Serret frame as intrinsic coordinates (ϕ, z) (Fig. 1e and Video S5, Methods). Interestingly, by following the orientation of this reference system along the midline, we noticed torsion associated with the cardiac looping, which is most pronounced around the AVC^14,15^. Plotting the intrinsic coordinates of the curved cylinder allows straightening and untwisting by implicitly removing the morphological torsion. Neglecting the actual distance to the midline and plotting the position along the midline (τ) against the angle (ϕ), we obtained a 2D projection (Fig. 1e, f and Video S5, Methods), in which the two outer curvature regions of atrium and ventricle are located side-by-side. A clear asymmetry in conduction speeds between cells at the inner and the outer curvatures was apparent in this representation (Fig. 1f). Irrespective of local bulging, however, the activation wave travelled smoothly in a ring-like fashion along the heart, as indicated by the isochronal lines in the cylindrical projection (Supplementary Fig. 7).

In order to document how the observed heterogeneity in cardiac function arises during cardiac looping, we extended our 3D optical mapping towards earlier developmental stages. During the crucial period between 36 and 52 hpf, ventricular cell number increased by about 45%, the initial heart tube developed into a two-chambered organ, the midline of the heart became increasingly curved and twisted (Supplementary Fig. 8), and the activation frequency increased – all signs of organ maturation. In spite of a net increase in cell numbers, the time required for activation to propagate from inflow to outflow decreased. A crowding of calcium transient activation dynamics indicated maturation of cells, with two groups differentiating from the early homogeneous pattern: working cardiomyocytes in atrium and ventricle (Fig. 2a-f). At 36 hpf, activation propagated evenly across the myocardium, compatible with peristalsis. Subsequently, calcium dynamics became increasingly structured. By 52 hpf, propagation of activation was fast across atrium and ventricle, while it remained slow in the AVC (Supplementary Fig. 9a). With organ maturation, cells in the ventricular part of the network showed longer calcium transient rise time (Supplementary Fig. 9b) but faster inter-cellular spread of activation (Fig. 2g-l, Supplementary Fig. 9c). Conduction also changed within chambers, such as along the outer curvature in the atrium, during these 16 hours. Increasing deformation and twisting of the cardiac tissue was associated with changes in cell shape (cf. Fig. 2d-f) and in conduction (Supplementary Fig. 10).

## Discussion

By noninvasively reconstructing the maturation process of the myocardium in its entirety at cellular resolution, our approach offers an integrative perspective on tissue and cell levels simultaneously. We show functional maturation in line with structural patterning of the heart muscle during development: starting from similar initial states, functionally distinct characteristics of calcium transients and conduction properties develop with a highly reproducible pattern relative to the cell locations (cf. Supplementary Fig. 7). Myocardial cells in the chambers remodel and specialize into functional tissue of working atrial and ventricular cardiomyocytes, while cells in the pacemaker and AVC regions continue to resemble the earlier phenotype from the tubular stage.

We demonstrate that myocardial activity can be recorded and analysed with cellular detail across the entire embryonic heart. Future technological advancements can extend the scope of our approach: First, genetically expressed voltage reporters with improved dynamic range can provide a closer readout of myocardial electrical activity. Second, cameras with higher speed and sensitivity would enhance the recording frequency of rapid volume scanning, needed to explore aberrant myocardial activation during arrhythmias. Third, the addition of optically gated actuators, such as light-activated ion channels or photo-pharmacological probes, would enable contact-free stimulation to probe the roles of individual cells or groups of cells in pacemaking, conduction, and arrhythmogenesis.

Our work further highlights the value of the zebrafish as a vertebrate model system for *in vivo* cardiology, especially when combined with high-speed light sheet microscopy and suitable data analysis pipelines. It opens the way to systematic, scale-bridging, *in vivo* studies of organogenesis by facilitating cell-accurate measurements across entire organs.

## Materials and Methods

### Fish husbandry and lines

Zebrafish (*Danio rerio*) were kept at 28.5 °C and handled according to established protocols^16^ and in accordance with EU directive 2011/63/EU as well as the German Animal Welfare Act. Transgenic zebrafish lines *Tg(myl7:GCaMP5G-Arch(D95N))^3^* and *Tg(myl7:H2A-mCherry)*^17^ were used. During the 1-cell stage, embryos were injected with morpholinos against tnnt2a to uncouple electrical and mechanical activity^18^.

### Sample preparation

Before imaging, fluorescent embryos were selected for absence of cardiac malformations and contractions, using an Olympus stereomicroscope equipped with an LED for transmitted light microscopy and a metal-halide light source and filter sets that match the excitation and emission spectra of GCaMP5G and mCherry for fluorescence excitation. Embryos were mounted in either 0.1 or 1.5% low gelling temperature agarose (Sigma A9414) inside cleaned polymer tubes (FEP tubing, inner/outer diameter 0.8/1.6 mm, BOLA S1815-04).

### Light sheet microscopy

We built a light sheet microscope for *in vivo* cardiac imaging in zebrafish embryos, based on a previously published design^7^. Imaging was performed in live zebrafish embryos between 36 and 52 hours post-fertilization (hpf) at a temperature of 24 °C. Heart rate at this temperature is 2 Hz, about 0.5 Hz lower than at the temperature recommended for breeding of 28.5 °C^19,20^. Embryos were kept in a custom imaging chamber filled with E3 fish medium and illuminated with a static light sheet generated from Coherent Sapphire LP lasers (488 and 561 nm) using a cylindrical lens and a Zeiss 10x/0.2 air illumination objective. Laser power was kept at or below 2 mW in the field of view (measured at the back aperture of the illumination objective) to exclude thermal effects on heart rate (an increase in heart rate was detected at laser powers of 5 mW and above). Fluorescence was collected and recorded using a Zeiss W Plan-Apochromat 20x/1.0 objective, a Zeiss 0.63x camera adapter, a Hamamatsu W-View image splitter and a Hamamatsu Flash 4.0 v2 sCMOS camera. Embryos were held in place by a Zeiss Lightsheet Z.1 sample holder and oriented using motorized translation and rotation stages (Physik Instrumente GmbH, Karlsruhe, Germany). For imaging of GCaMP5G, a z-stack of movies covering the entire heart was recorded by moving each embryo through the light sheet (488 nm excitation, band-pass 525/50 nm emission filter, 2.5 ms exposure time = 400 Hz, 1 μm z-steps). For imaging of H2A-mCherry, a matching z-stack was recorded immediately afterwards (561 nm excitation, long-pass 561 nm emission filter, 20 ms exposure time = 50 Hz, 1 μm z-steps). Image acquisition was controlled by a custom program written in LabView (National Instruments). Images were streamed onto a RAID-0 array of four 512 GB solid-state drives.

### Data analysis

#### Segmentation of cell nuclei

Volumes of cell nuclei were extracted from the H2A channel of the raw image volumes using grayscale blob detection^21^. To resolve potential errors in automated processing (false detection, missing cells), cell positions were manually curated using an in-house software facilitating evaluation, addition and deletion of nuclear positions in the 3D data sets. Curated nuclei positions were used for further processing.

#### Signal extraction and processing

From the measured nuclear positions, a reference volume was extracted for each cell by computing the Euclidean distance transform and thresholding the distance field around each centroid (nucleus) at a radius of 5 voxels. For each cell a calcium transient was extracted at each time step by averaging the signal over all voxel within the reference volume. The resulting transients were processed using a low pass filter (with a cutoff frequency of 5 Hz) to reduce noise, and finally normalized to the range [0,1].

#### Extraction of midline

The heart’s midline was manually traced using an in-house software. The user could draw the midline in two different 2D projections (front and top view) by placing discrete points that were connected across the different projections. The final midline was then obtained by a 4^th^ order B-Spline interpolation to facilitate subsequent computation of intrinsic geometric characteristics such as curvature and torsion.

#### Frenet-Serret frame

To describe each cell’s position within the myocardium, we established a curved cylindrical coordinate system. The position along the midline (τ) was calculated for each cell by finding the closest point on the midline for the respective centroid. The moving coordinate system along the midline was then computed using the Frenet-Serret frame ^22^, calculating the local tangent (***T***), normal (***N***), and binormal (***B***) vectors:

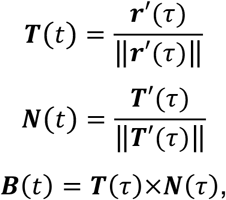

with being the actual 3D position vector 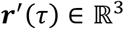 on the midline parameterized by τ. Then each cell’s position was mapped to this new coordinate system as (τ, ϕ, z), where τ was the position along the midline (normalized to 0 and 1 after reparameterization), ϕ the angle with respect to the local binormal vector (in the local **N-B** plane), and z the radial distance of the centroid from the midline.

#### Cylindrical 2D map projections

To map cell positions from 3D to 2D, we projected the curved cylindrical coordinates to the first two components (τ, ϕ, *z*) → (*τ*, ϕ), discarding the radial distance from the midline. This projection facilitates global visualization of the myocardium, irrespective of its looping stage, in a single flat projection by cutting the cylinder at ϕ = ±*π* and plotting the coordinates in a rectangular coordinate system in the range *τ* ∈ [0, 1] and ϕ ∈ [−*π*, *π*], respectively.

#### Topology reconstruction

The myocardial topology (local connectivity of cells) was estimated by projecting the 3D centroid positions of each cell and its nearest neighbors locally into the 2D Euclidean plane using Singular Value Decomposition^23^ of the 3D centroid positions. In each local 2D projection, the cellular connectivity was estimated by computing the 2D Delaunay triangulation^24^. The estimated edges *e_ij_* between cells and the original 3D centroid positions of each cell *v_i_* were used to abstractly represent the myocardium as an undirected graph *G* = (*V*, *E*) where *v_i_* ∈ *V*, *e_ij_* ∈ *E*.

#### Computation of biological conduction speed

As the spread of activation in cardiac tissue is governed by cell activation and time delay at gap junctions between cells, we used the graph representation to estimate the average local cell-to-cell conduction speed *cs_i_* in terms of (dimensionless) links traversed per unit of time. Thus, at each cell’s position the (harmonic) mean of the number of traversed edges per time was computed over all paths from the cell to its neighbors: 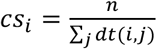 for all neighbors *j* of cell *i* with *dt*(*i,j*) being the temporal difference in activation time between cell *i* and cell *j*.

#### Estimation of cell size and shape

We further used the graph representation to extract an estimate of a cell’s shape by computing the principal components of the distribution of neighboring nuclei around each cell. Thus, each cell is represented as an ellipsoidal region defined by the eigenvectors *ν_i_* and eigenvalues *λ_i_* of the local principal value decomposition. Cell shape was then approximated as the fractional anisotropy of each ellipsoidal region: 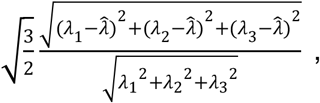 with 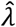 being the average eigenvalue. Cell size was approximated as the volume 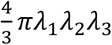 of each ellipsoid.

#### Pacemaker identification

Potential pacemaker cells were defined as the cells that showed earliest activation (activation time shorter than the 0.05-quantile of overall distribution of activation times) and that conducted slowly (conduction speed slower than the median conduction speed within the cell population). All cells falling within this category were labeled as potential pacemaker cells.

## Acknowledgments

We thank the entire Huisken Lab, in particular R. Power and M. Mickoleit, as well as A. El-Armouche, V.M. Christoffels, D.J. Christini, F. Ortega, L. Herzel and K. Thierbach for valuable discussions. N.S. and J.H. are supported by the ERC Consolidator Grant SmartMic and P.K. is supported by the ERC Advanced Grant CardioNECT.

## Author contributions

M.W. built the instrument and performed the experiments, N.S. analyzed and visualized the data, all authors contributed to conceiving the work, interpreting data, and writing of the manuscript.

## Supplementary Figures

**Supplementary Fig. 1:**
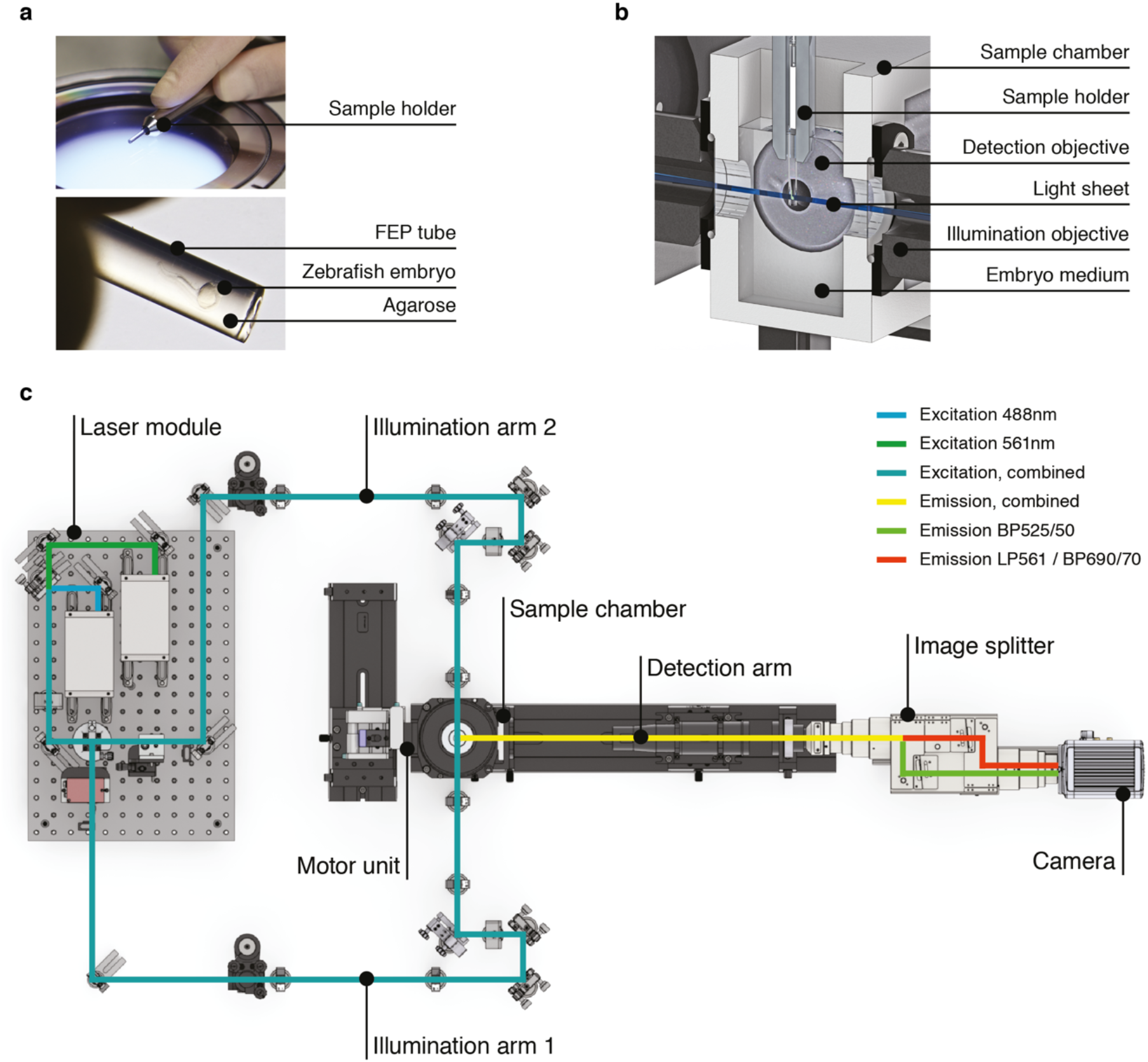
High-speed light sheet microscopy for *in vivo* 3D optical mapping. **(A)** A zebrafish embryo is mounted in agarose inside a fluorinated ethylene propylene (FEP) tube. (**B**) Section view of the sample holder with mounted zebrafish embryo placed inside the medium-filled sample chamber. The embryo is placed in the field of view of the detection objective and illuminated with a static light sheet from one of two sides. (**C**) Top view of the high-speed light sheet microscope for *in vivo* cardiac imaging. The laser module combines a 488 and a 561nm laser line and sends the beam into the two illumination arms. Both arms generate identical light sheets from two opposite sides. The motor unit positions the sample holder with the mounted zebrafish embryo at the intersection of illumination and detection path. Fluorescence emission is split and recorded with an sCMOS camera running at up to 400Hz.

**Supplementary Fig. 2:**
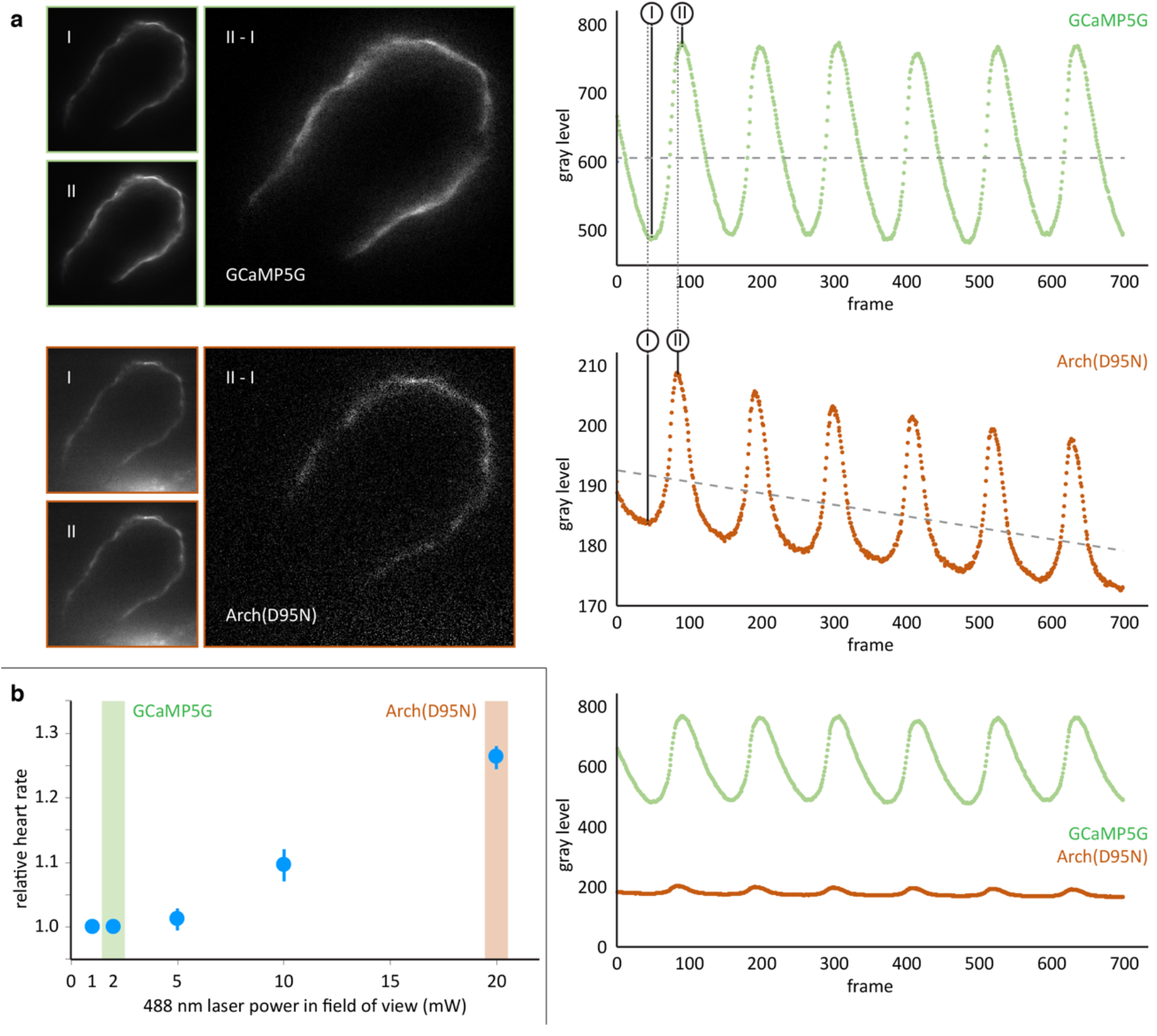
Comparison of the calcium reporter GCaMP5G and the voltage reporter Arch(D95N) for multi-scale readout of cardiomyocyte activation. **(a)** Optical section across the atrium of a zebrafish embryo at 52 hpf expressing GCaMP5G and Arch(D95N) in cardiomyocytes. Both channels are recorded simultaneously. Smaller images: raw data recorded at lowest (I) and highest (II) fluorescence signal, as indicated in the intensity plots. Note how intensity plots illustrate the known slight delay between intensity maxima of calcium versus voltage traces, and overall excellent capture of presence and temporal dynamics of electrical activation. Larger images: results of image I subtracted from image II, presenting the maximum intensity difference (image brightness adjusted independently for better visibility). Plots show mean raw intensities over time measured along the myocardium visible in the images. **(b)** Laser powers in the field of view (measured at the back pupil of the illumination objective) used for the experiment presented in (a) and their impact on the heart rate (n = 3 zebrafish embryos). Arch(D95N) required 10x more laser power than GCaMP5G, yielded a very low signal with about 20 gray levels dynamic range across one cardiac cycle, and was affected by bleaching. Zebrafish embryos showed increased heart rate when illuminated with the high laser power needed for Arch(D95N) imaging. GCaMP5G showed a dynamic range of about 300 gray levels at an order of magnitude lower laser power, with no signs of visible photobleaching or illumination-induced increase in heart rate.

**Supplementary Fig. 3:**
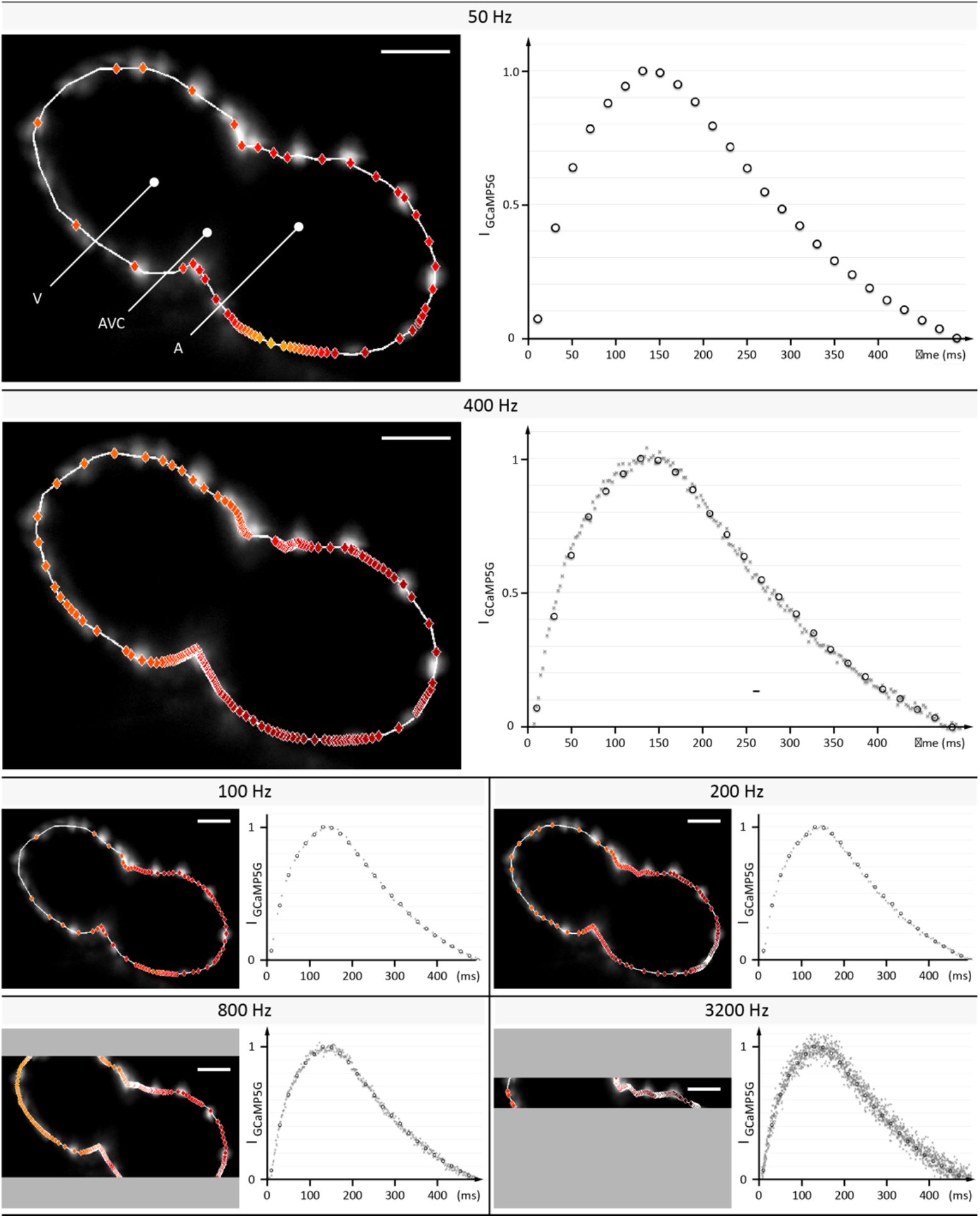
Evaluation of signal coverage and tracing precision using the fluorescent calcium reporter GCaMP5G at different recording speeds. Optical sections across the silenced heart of a 2 days post fertilization (dpf) zebrafish embryo expressing GCaMP5G and H2A-mCherry in cardiomyocytes, recorded at 50 to 3,200 Hz (exposure times 20 to 0.3 ms) with a constant pixel size of 0.5 μm. Raw image data of H2A-mCherry is shown to mark cell positions. Red markers indicate location of peak fluorescence intensities across a single cardiac cycle. Gray areas in 800 Hz and 3,200 Hz images indicate the proportion of the field of view that could not be recorded, as the number of lines imaged is the speed-limiting factor on the sCMOS camera. Plots show normalized fluorescence intensity over time, measured in a sub region with GCaMP5G signal. Circles show data points from the measurement at 50 Hz as reference. Higher recording speed results in a better representation of the calcium transient until 400-800 Hz, especially during the initial rise in intensity. At an even higher rate of 3,200 Hz, noise deteriorates the signal. Peak intensities could be traced with cellular precision at 400 and 800 Hz, before signal noise reduced precision.

**Supplementary Fig. 4:**
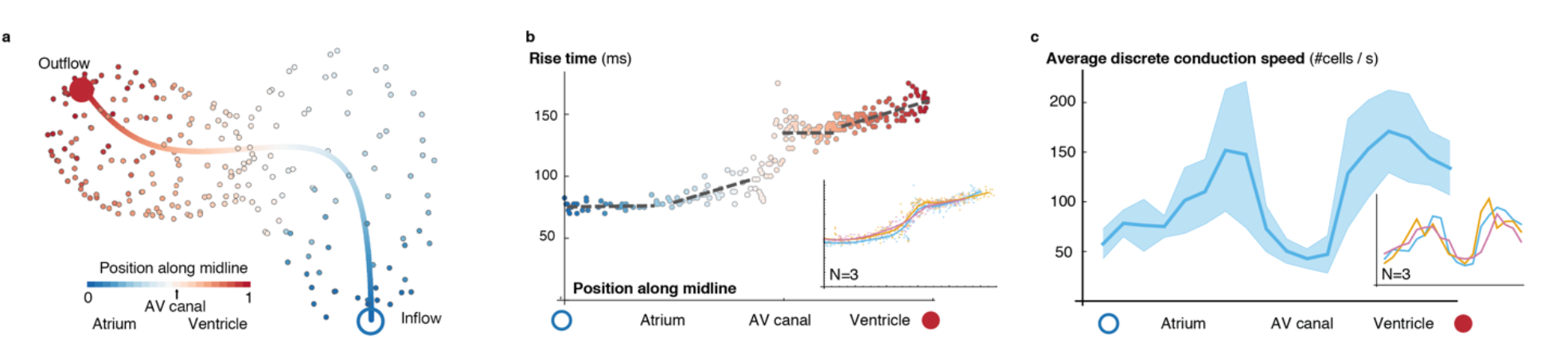
Activation and conduction properties of cells vary from inflow to outflow, with patterns conserved across different hearts (52 hpf, n=3). **(a)** The centerline is traced in 3D from inflow (blue) to outflow (red), providing the base for a canonical ordering of the cells. Color code indicates position of cell along midline. (**b**) Populations of cells with different rise times can be distinguished close to the inflow tract, in the atrium and in the ventricle (indicated by different linear fits), with a pronounced discontinuity in the atrio-ventricular canal (AVC). Color code indicates position of cell along midline. (c) Distribution of biological conduction speeds across the heart, from inflow (blue circle) to outflow (red disk), mean and standard deviation. The inset shows the mean conduction speed for three different hearts at 52 hpf.

**Supplementary Fig. 5:**
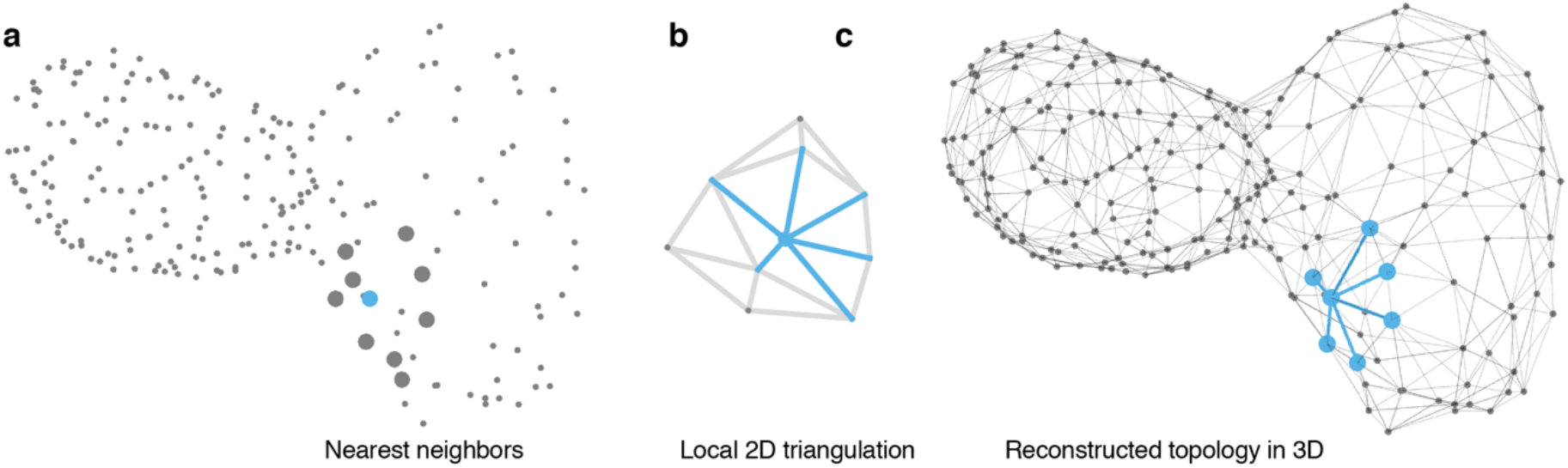
Reconstruction of myocardial topology. **(a)** Cell positions are indicated by points at the optically identified location of their nucleus in 3D. Potential neighbors for each cell are identified by taking nearest neighbors with respect to their Euclidean distance in 3D. A sample cell is highlighted in blue and the respective nearest neighbors by large gray dots. **(b)** The centroids are locally projected into a 2D plane and the Delaunay triangulation (gray and blue lines) is computed to extract the topological connectivity between cells. Edges connecting the reference cell with its neighbors in the 2D projection are highlighted in blue. **(c)** The connections between the reference cell and its neighbors (blue) are projected back into the original 3D space. Iterating this procedure for all cells yields the final topology (gray edges).

**Supplementary Fig. 6:**
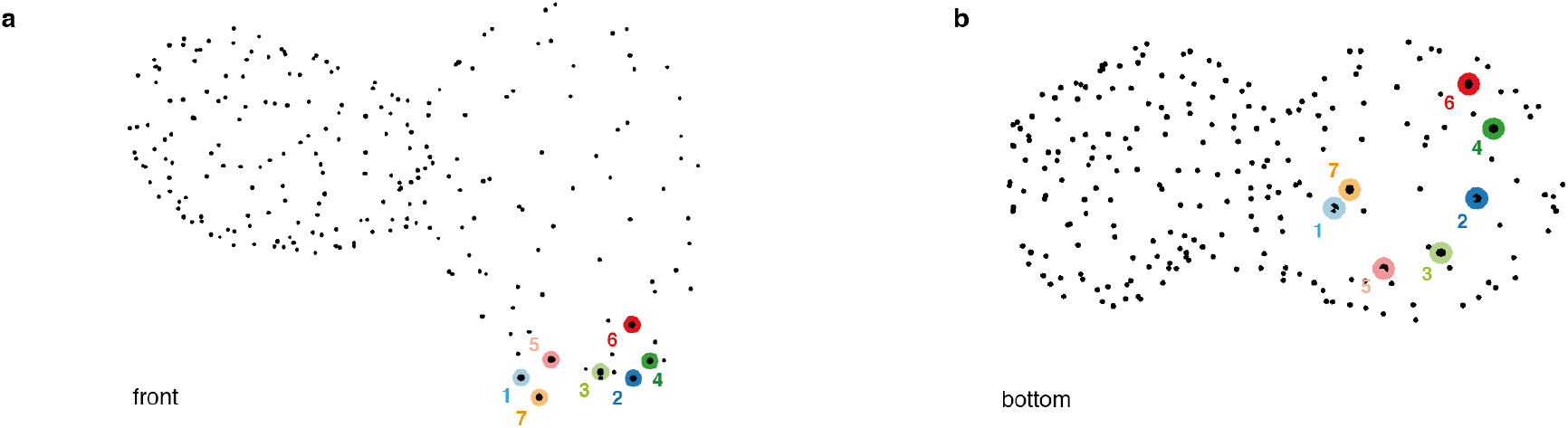
Location of pacemaker cells. **(a, b)** Pacemaker cells (shown as colored disks), all located in a ring-like region at the atrial inflow region, shown in front (a) and bottom view (b).

**Supplementary Fig. 7:**
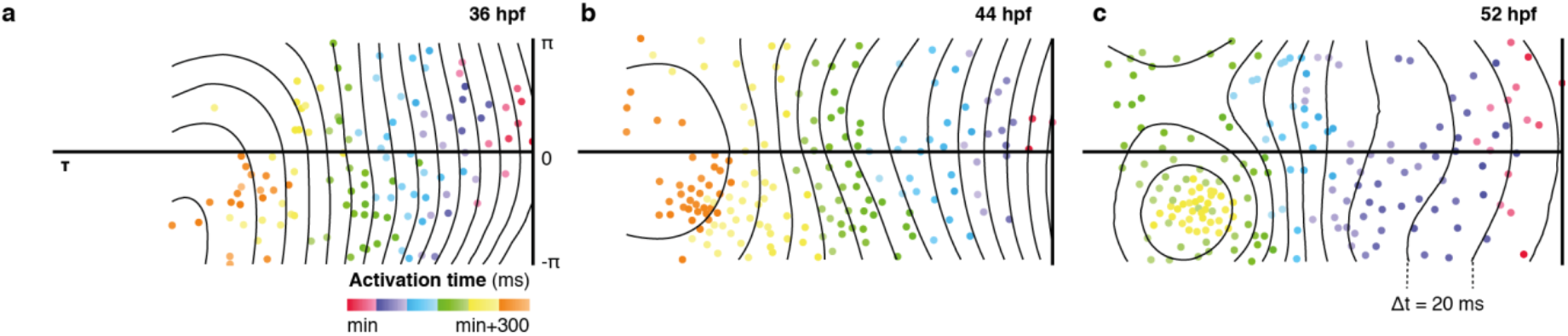
Patterns of cell activation across myocardium. Global distribution of cellular activation times shown as 2D projections, illustrate planar activation in spite of increasingly complex chamber morphology and matching heterogeneity in individual cell properties. Isochronal lines shown in black (Δt = 20 ms).

**Supplementary Fig. 8:**
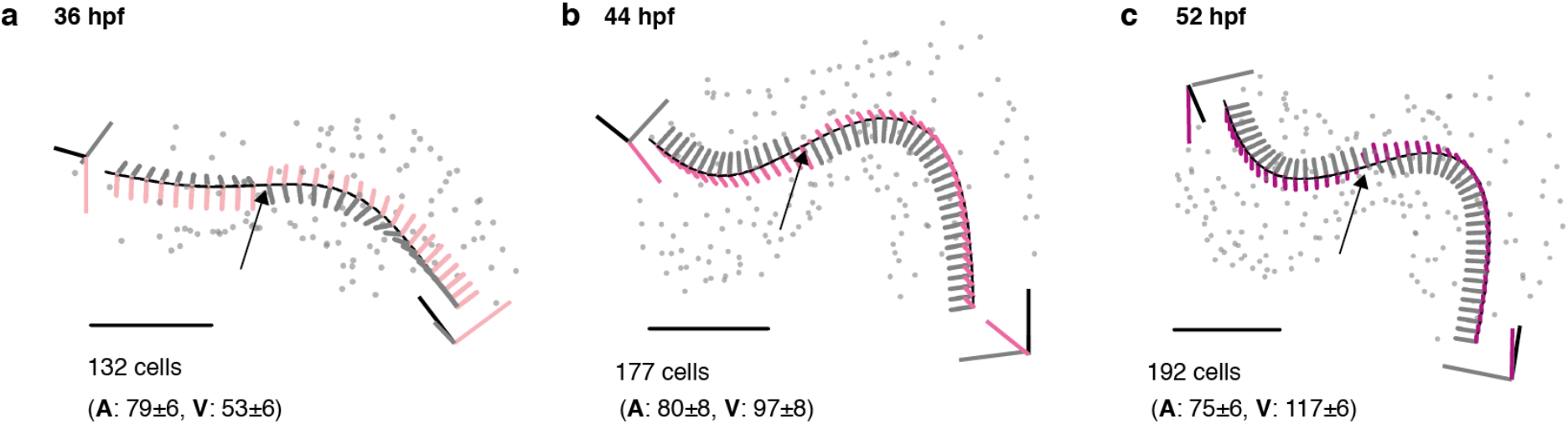
Structural organ maturation from 36 to 52 hpf. (**a**) One and the same developing heart, imaged at three different time-points (36 (a), 44 (b), and 52 hpf (c), scale bar 50 μm. The fitted centerline visualizes the re-shaping of the myocardium during the ongoing looping / twisting process in early cardiac development. Local coordinate systems are shown along the midline (tangent – black, normal – grey, binormal – color), the twisting point is highlighted by an arrow. Total and chamber-specific cell numbers are indicated for each stage of development (A - atrial cells, V - ventricular cells).

**Supplementary Fig. 9:**
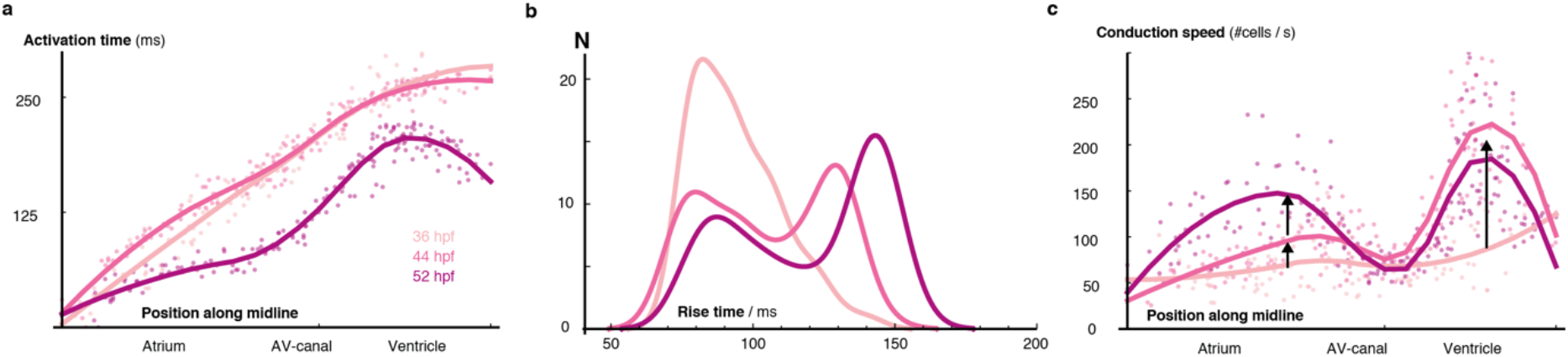
Developmental changes of cellular characteristics along the heart. (**a**) The distribution of activation times is shown for three time points: 36, 44 and 52 hpf. Dots indicate individual cell activation times. Smooth regression profiles are depicted as solid lines and show faster activation with progressive organogenesis. (**b**) The distribution of rise times during development. The probability density function of each developmental stage is shown by a smooth kernel density. The ordinate axis shows cell numbers and illustrates progressive emergence of cell sub-populations with different calcium transient properties. (**c**) The distribution of each cell’s conduction speed along the midline is shown as dots. Solid lines indicate smooth regression profiles for each stage. All hearts where normalized to a standard length, showing how speed of biological conduction in working myocardium rises by comparison with early developmental stages, whereas AVC speed is unchanged.

**Supplementary Fig. 10:**
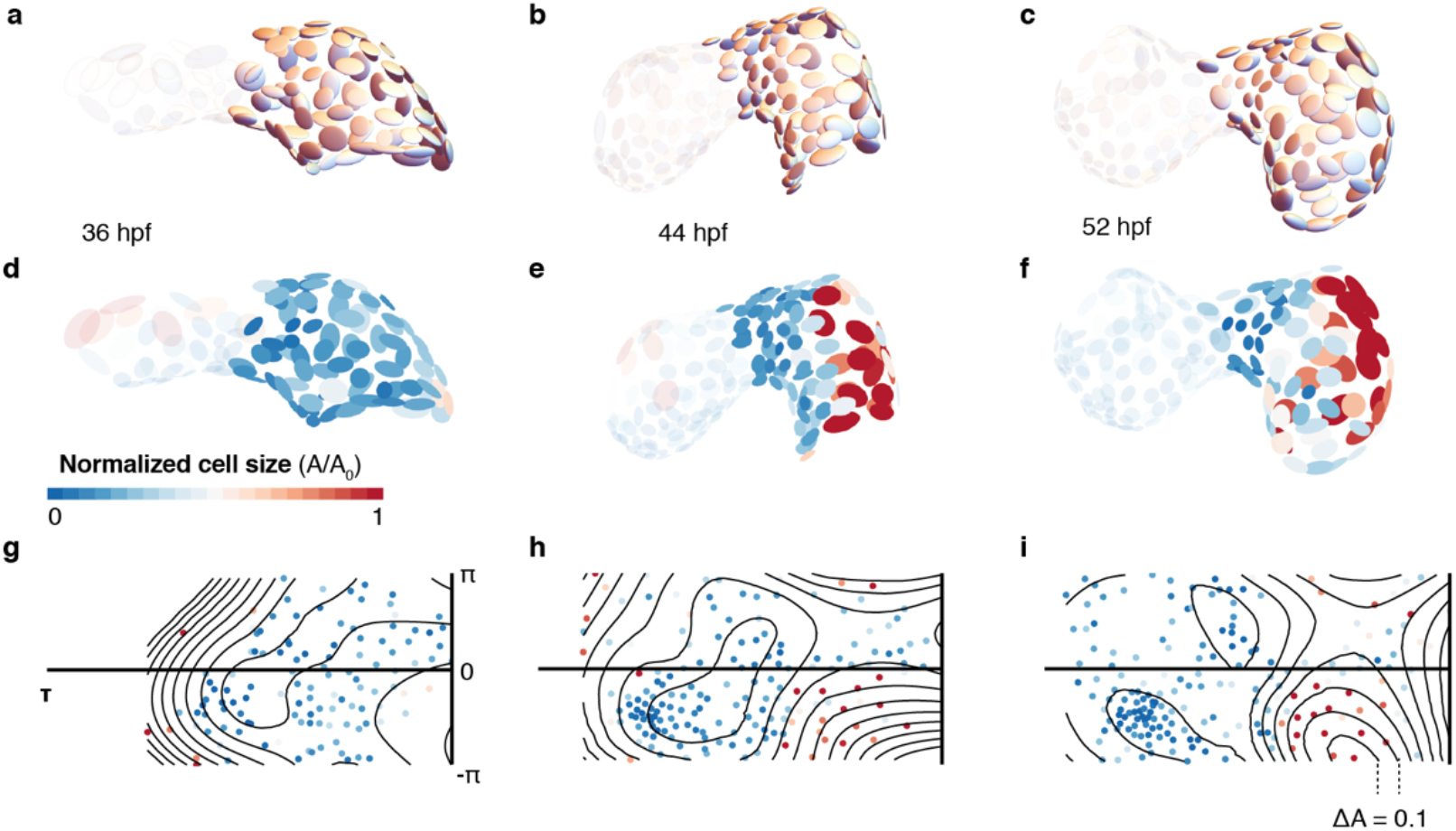
Cell shape changes during development. Cell shapes in the atrium are represented by scaled ellipsoids for the same heart at 36 hpf (**a**), 44 hpf (**b**), and 52 hpf (**c**). Ventricular cells are shown with reduced opacity. (**d, e, f**) The same cells as in (**a, b, c**) with additional color code indicating cell size. Smaller cells are shown in blue, larger cells in red, highlighting the increase in cell size, in particular along the larger curvature of the atrium. Changes in cell size shown as 2D projections for 36 (**g**), 44 (**h**), and 52 hpf (i). The color code indicates normalized cell size as in (d, e, f). Isosize lines shown in black (ΔA = 0.1).

## Supplementary Video Legends

**Video S1: Raw GCaMP5G signal at different imaging depths**

Fluorescence signal recorded at 400 Hz at three different depths (50, 90 and 140 μm along the optical axis) in the heart of a living Tg(myl7:GCaMP5G) zebrafish at 52 hpf using the described imaging setup.

**Video S2: Calcium signal at different imaging depths after synchronization**

Raw data from Video S1 after completed post-acquisition synchronization.

**Video S3: 4D reconstruction of cellular calcium transients**

(1) 4D reconstruction: Reconstruction of calcium activation from the synchronized planar movies recorded in the heart of a living Tg(myl7:GCaMP5G) zebrafish embryo at 52 hpf. The number of slices used in the reconstruction is indicated by n. Raw GCaMP5G signal is shown in orange. Time is indicated in ms.

(2) Cell detection: The raw fluorescence signal of myl7:H2A-mCherry is overlaid with the centroids of detected nuclei.

(3) Calcium mapping: The raw myl7:GCaMP5G signal is shown. The dots indicate the extracted cellular positions.

(4) Cell specific transients: The normalized GCaMP5G transients are shown for two sample cells (one atrial and one ventricular cell).

**Video S4: Computation of biological conduction speed**

(1) Activation timing: Cell positions are indicated by gray dots. Activated cells are highlighted in yellow. Time is shown in ms.

(2) Network reconstruction: Estimated local topology is shown as gray edges connecting neighboring cells.

(3) Activation across network: Activated cells in the network are highlighted in yellow.

(4) Cell-cell conduction speed: The average conduction speed is shown for each cell after activation of its neighbors in the network. Color code indicates conduction speed (red - high, blue - low).

**Video S5: Mapping of myocardial geometry.**

(1) Tracing of midline: The black line traces the center of the myocardium from inflow to outflow. Cellular positions are shown as gray spheres.

(2) Moving reference frame: The intrinsic reference frame is shown along the extracted midline. The three axes represent the tangent (black), normal (gray), and binormal (blue) vectors. The trace of the binormal vector is shown as blue band behind the moving trihedron.

(3) Untwisting: The heart is successively untwisted by reducing the torsion of the midline to 0. The gray lines indicate traces of cells during this process.

(4) Straightening: The heart is straightened by reducing the remaining curvature of the midline to 0.

(5) Projection to cylinder. Each cell is projected to the same radial distance from the straightened midline resulting in a cylindrical reference system.

(6) Unrolling: The cylinder is unfolded into the 2D plane, resulting in the 2D plots used in the main text. Color code in the final frame indicates biological conduction speed.

## Supplementary Information

Videos S1-S5

## Reference

1 Chi, N. C. et al. Genetic and physiologic dissection of the vertebrate cardiac conduction system. PLoS Biol. 6, e109 (2008).

2 Scherz, P. J., Huisken, J., Sahai-Hernandez, P. & Stainier, D. Y. R. High-speed imaging of developing heart valves reveals interplay of morphogenesis and function. Development 135, 1179–1187 (2008).

3 Mickoleit, M. et al. High-resolution reconstruction of the beating zebrafish heart. Nat. Methods 11, 919–922 (2014).

4 Bouchard, M. B. et al. Swept confocally-aligned planar excitation (SCAPE) microscopy for high-speed volumetric imaging of behaving organisms. Nat. Photonics 9, 113–119 (2015).

5 Fahrbach, F. O., Voigt, F. F., Schmid, B., Helmchen, F. & Huisken, J. Rapid 3D light-sheet microscopy with a tunable lens. Opt. Express 21, 21010–21026 (2013).

6 Hou, J. H., Kralj, J. M., Douglass, A. D., Engert, F. & Cohen, A. E. Simultaneous mapping of membrane voltage and calcium in zebrafish heart in vivo reveals chamber-specific developmental transitions in ionic currents. Front. Physiol. 5, 344 (2014).

7 Liebling, M., Forouhar, A. S., Gharib, M., Fraser, S. E. & Dickinson, M. E. Fourdimensional cardiac imaging in living embryos via postacquisition synchronization of nongated slice sequences. J. Biomed. Opt. 10, 054001 (2005).

8 Kralj, J. M., Douglass, A. D., Hochbaum, D. R., Maclaurin, D. & Cohen, A. E. Optical recording of action potentials in mammalian neurons using a microbial rhodopsin. Nat. Methods 9, 90–95 (2012).

9 Brandenburg, S. et al. Axial tubule junctions control rapid calcium signaling in atria. J. Clin. Invest. 126, 3999–4015 (2016).

10 Moorman, A. F., de Jong, F., Denyn, M. M. & Lamers, W. H. Development of the cardiac conduction system. Circ. Res. 82, 629–644 (1998).

11 Arrenberg, A. B., Stainier, D. Y. R., Baier, H. & Huisken, J. Optogenetic control of cardiac function. Science 330, 971–974 (2010).

12 Poon, K. L. & Brand, T. The zebrafish model system in cardiovascular research: A tiny fish with mighty prospects. Glob Cardiol Sci Pract 2013, 9–28 (2013).

13 Auman, H. J. et al. Functional modulation of cardiac form through regionally confined cell shape changes. PLoS Biol. 5, e53 (2007).

14 Harvey, R. P. Patterning the vertebrate heart. Nat. Rev. Genet. 3, 544–556 (2002).

15 Männer, J. Cardiac looping in the chick embryo: a morphological review with special reference to terminological and biomechanical aspects of the looping process. Anat. Rec. 259, 248–262 (2000).

16 Nusslein-Volhard, C. & Dahm, R. Zebrafish: A practical approach. (Oxford University Press, 2002).

17 Schumacher, J. A., Bloomekatz, J., Garavito-Aguilar, Z. V. & Yelon, D. tal1 Regulates the formation of intercellular junctions and the maintenance of identity in the endocardium. Dev. Biol. 383, 214–226 (2013).

18 Sehnert, A. J. et al. Cardiac troponin T is essential in sarcomere assembly and cardiac contractility. Nat. Genet. 31, 106–110 (2002).

19 Baker, K., Warren, K. S., Yellen, G. & Fishman, M. C. Defective ‘pacemaker’ current (Ih) in a zebrafish mutant with a slow heart rate. Proc. Natl. Acad. Sci. U. S. A. 94, 4554–4559 (1997).

20 Kopp, R., Schwerte, T. & Pelster, B. Cardiac performance in the zebrafish breakdance mutant. J. Exp. Biol. 208, 2123–2134 (2005).

21 Lindeberg, T. Feature Detection with Automatic Scale Selection. Int. J. Comput. Vis. 30, 79–116 (1998).

22 Kreyszig, E. Differential Geometry. (Courier Corporation, 1959).

23 Bronshtein, I. N. Handbook of Mathematics. (Springer, 2007).

24 Delaunay, B. N. Sur la Sphère Vide. Biol. Bull. Acad. Sci. USSR 793–800 (1934).

